# EEG Movement Tagging Objectively Measures Biological Motion Perception

**DOI:** 10.1101/2022.05.03.490439

**Authors:** Emiel Cracco, Danna Oomen, Liuba Papeo, Jan R. Wiersema

## Abstract

Detecting biological motion is essential for adaptive social behavior. Previous research has revealed the brain processes underlying this ability. However, brain activity during biological motion perception captures a multitude of components. As a result, it is often unclear which components reflect movement processing and which components reflect secondary processes building on movement processing. To address this issue, we developed a new approach that objectively defines the brain response associated with biological motion perception.

Specifically, we showed 30 male and female adults a point-light walker moving at a pace of 2.4 Hz and used EEG frequency tagging to measure the brain response coupled to that pace (‘movement tagging’). The results revealed a reliable response at the walking frequency that was reduced by two manipulations known to disrupt biological motion perception: phase scrambling and inversion. Interestingly, we also identified a brain response at half the walking frequency (i.e., 1.2 Hz), corresponding to the rate at which the individual dots completed a cycle. In contrast to the 2.4 Hz response, the response at 1.2 Hz was increased for scrambled walkers. These results show how frequency tagging can be used to objectively measure the visual processing of biological movements and can dissociate between global (2.4 Hz) and local (1.2 Hz) processes involved in biological motion perception, at different frequencies of the brain signal.

---

The ability to recognize other living beings is key for any organism living in a social environment. One of the most important indicators of ‘life’ is biological motion (Troje & Westhoff, 2006). Perhaps for this reason, humans as well as other species have developed dedicated mechanisms to detect biological movement (Grossman et al., 2005; Pitcher & Ungerleider, 2021). These mechanisms typically make use of both ‘shape’ and ‘motion’ cues (Giese & Poggio, 2003; Hirai & Senju, 2020; Lange & Lappe, 2006), but can also detect biological movement from motion cues alone (Giese & Poggio, 2003). To study the role of motion cues, research often uses ‘point-light figures’, a type of stimuli that depict human (or animal) movements as a set of moving dots placed on the major joints of the body (Blake & Shiffrar, 2007; Johansson, 1973). More specifically, because these figures convey little more than kinematic information, observers have to make use of the local dot trajectories and the relationships between them to identify the global movement pattern in the stimulus (Giese & Poggio, 2003).

Several paradigms have been developed to study this process. In one of the most common paradigms, participants have to indicate whether a point-light figure is present in an array of noise made up from scrambled dots that maintain the same local movement trajectory as the original figure, but are placed in different spatial locations (Troje, 2013). Although this procedure makes it impossible to identify biological movement from local motion features, research has shown that point-light figures can still be detected reasonably well under such circumstances (Bertenthal & Pinto, 1994; Chang & Troje, 2009), except when they are presented upside-down (Bertenthal & Pinto, 1994; Pavlova & Sokolov, 2000). This suggests that the spatial relations between the different body parts are important to detect biological motion (Giese & Poggio, 2003; Lange & Lappe, 2006) and hence confirms that biological motion perception relies at least partly on global, configural processing (Maurer et al., 2002).

Another approach to study biological motion processing is to measure brain activity with fMRI or EEG while participants passively observe moving point-light figures. Studies using fMRI have shown that biological movement perception activates a wide network of visual brain areas, including the extrastriate and fusiform body areas, the medial temporal cortex, and the superior temporal sulcus. In line with behavioral evidence, activity in these brain areas is reduced by manipulations that perturb the global movement percept, such as scrambling (e.g., Engell & McCarthy, 2013; Grossman et al., 2000; Peuskens et al., 2005) and inversion (e.g., Grossman & Blake, 2001; Pavlova et al., 2017; Peuskens et al., 2005). EEG studies have instead focused mainly on the time course of event-related potentials (ERPs) elicited during biological motion perception (e.g., Chang et al., 2021; Hirai et al., 2003; White et al., 2014). The results suggest that scrambling has an effect already after 150-200 ms, whereas the effect of inversion emerges only after 400 ms (White et al., 2014). Thus, despite having similar effects, scrambling and inversion appear to operate at different processing stages.

However, a key challenge is that brain activity during biological motion perception captures not only movement processing but also higher-order processes building on it (e.g., White et al., 2014), such as the extraction of social features (e.g., emotional state, sex, identity, …) from movement kinematics (e.g., Atkinson et al., 2004; Cutting & Kozlowski, 1977; Kozlowski & Cutting, 1977; Pavlova, 2012). To address this issue, we propose to use frequency tagging (Norcia et al., 2015). More specifically, we propose to lock brain responses to a cyclical movement (e.g., walking) repeating at a fixed rate, an approach we will refer to as ‘movement tagging’. In case of walking, this should elicit a brain response that recurs every time a footstep is completed. As a result, the brain response is defined by the temporal profile of the movement and movement processing gets separated from other processes that also take place during movement perception, but are not specifically tied to the movement itself.

Hence, by eliciting cyclical brain responses coupled to the rate of movement repetition, frequency tagging can provide an objective measure of movement processing. Interestingly, however, while a number of studies have already combined frequency tagging with biological motion stimuli (Alp et al., 2017; Burton et al., 2016; Cracco et al., 2021; Hasan et al., 2017; Zarka et al., 2014), only one study tagged the movement itself (Cracco et al., 2021). That is, Cracco et al. (2021) presented repeating sequences of static body postures that either did or did not form a fluent apparent movement. The results revealed dissociable responses at frequencies linked to posture and movement presentation, suggesting that movement tagging can indeed be used to measure movement processing. However, by using apparent instead of actual motion, this study did not measure the visual processing of human movement, but rather its top-down reconstruction from sequences of static body postures, which is known to rely on fundamentally different processes (e.g., Giese & Poggio, 2003; Orgs et al., 2016).

In sum, the aim of this study was to measure the visual processing of human movement from kinematic information. To this end, we developed a novel approach that separates movement processing from other processes not directly tied to the movement. More specifically, we presented a point-light figure walking at a fixed frequency (2.4 Hz) and measured brain responses coupled to that frequency (Figure 1). At the same time, we also manipulated two variables known to perturb biological motion perception: phase scrambling (e.g., Beintema et al., 2006; Troje & Westhoff, 2006) and body inversion (e.g., Bertenthal & Pinto, 1994; Pavlova et al., 2017; White et al., 2014). If the measured brain response indeed reflects biological movement processing, it should be reduced when the walker is scrambled or inverted.

**Figure 1.**
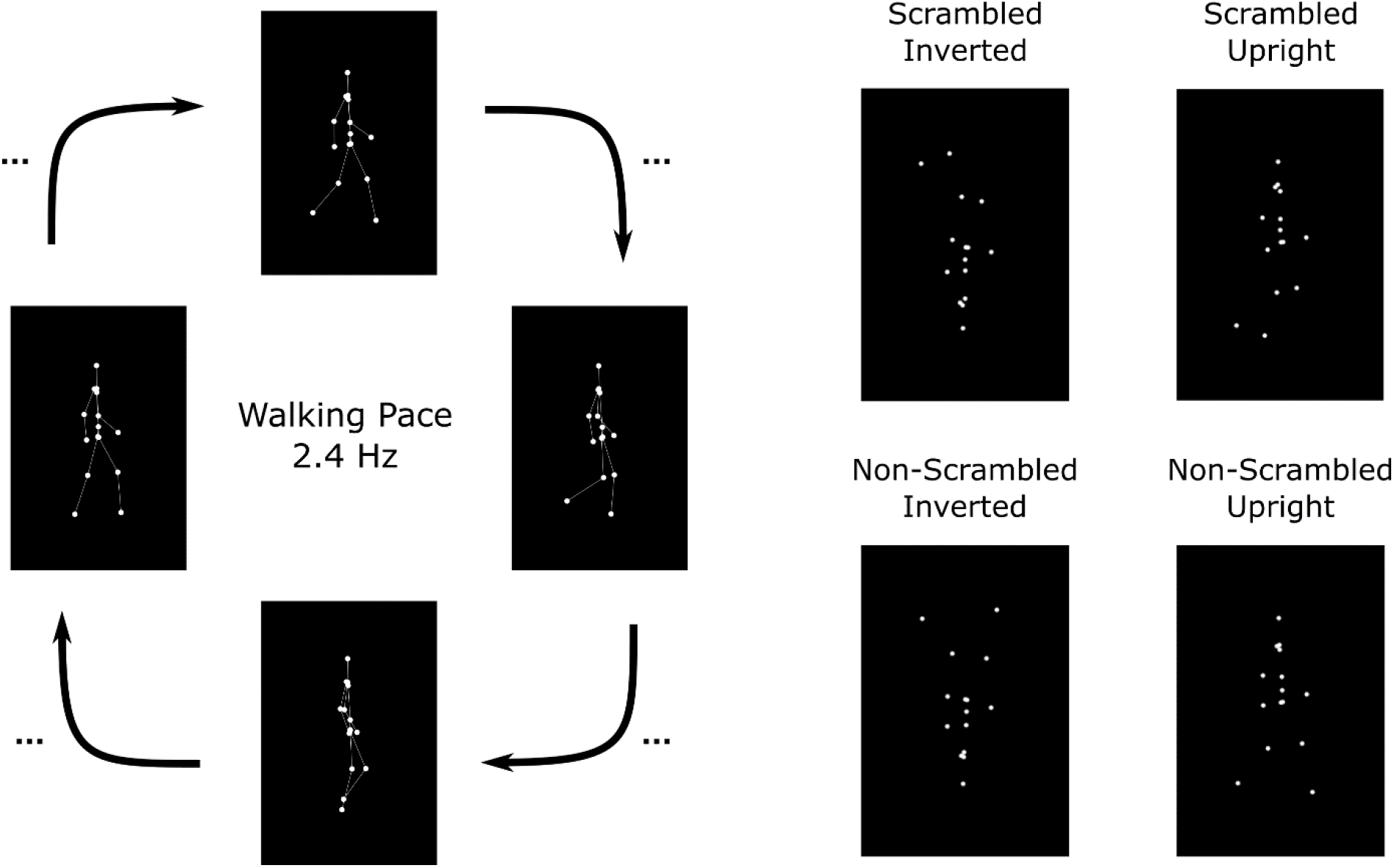
Experimental Task *Note*. In the task, participants saw a point-light figure walking at a fixed pace of 2.4 Hz. The walker could be presented scrambled and inverted, scrambled and upright, non-scrambled and inverted, or non-scrambled and upright. Lines between the dots were not shown in the actual experiment. Example videos are available on the Open Science Framework (OSF; https://osf.io/xwgmy/?view_only=b4491d96fc9e443e87417ec20f1d9afc).

## Open Science Statement

This study was preregistered (https://aspredicted.org/1P9_PNW). The stimuli, data files, analysis script, and experimental program will be made available on the OSF upon publication.

## Methods

### Participants

As this is the first study that uses frequency tagging to measure brain responses coupled to point-light movements, we did not have strong a-priori expectations regarding the effect size. However, given that frequency tagging is known for its high stimulus-to-noise ratio (Norcia et al., 2015) and that scrambling and inversion are known to reliably disrupt biological motion perception (e.g., Bertenthal & Pinto, 1994; Grossman et al., 2000; Grossman & Blake, 2001), we reasoned that small effect sizes were unlikely. Therefore, we decided to base our sample size on a power analysis assuming 80% power for a medium effect size of *d*_z_ = 0.50. This revealed that we needed a sample size of *N* = 33 to detect such an effect. We further preregistered that we would collect 3 more participants if *N* < 30 following exclusions, until *N* ≥ 30. Unfortunately, due to an undetected technical issue leading to bad data quality for a subset of participants (i.e., > 10% of electrodes requiring interpolation), 9 participants had to be excluded after collecting the first 33 participants. As preregistered, we therefore collected 6 (= 2 × 3) additional participants, leading to an eventual sample size of *N* = 30 (10 male, 20 female, *M*_age_ = 23.03, *range*_age_ = 18–33). All participants had normal or corrected-to-normal vision, had no history of neurological or psychiatric disorder, and signed an informed consent before the experiment. All procedures were approved by the ethics board of the Faculty of Psychological and Educational Sciences at Ghent University (2021/129).

### Task, Stimuli, and Procedure

During EEG preparation, participants first filled out the Dutch version of the Autism Quotient questionnaire (AQ; Hoekstra et al., 2008). The AQ had good internal consistency in the current study (α = 0.78) and was included for exploratory purposes, based on previous research suggesting anomalous processing of biological motion in autism (Federici et al., 2020; Todorova et al., 2019; Van der Hallen et al., 2019). Afterwards, during the experiment, participants were seated in a Faraday cage, approximately 80-100 cm from a 24-inch computer monitor with a 60 Hz refresh rate. The experiment was programmed in PsychoPy (Peirce et al., 2019) and consisted of 16 blocks in which participants observed a point-light walker moving at a fixed frequency of 2.4 Hz (i.e., 1 step every ∼ 417 ms) for a duration of 128 steps (∼ 53 sec), including a 4-step fade-in and a 4-step fade-out period (∼ 2 sec each). All point-light figures were created using the online BMLStimuli tool (Troje, 2002, 2008; https://www.biomotionlab.ca/Experiments/BMLstimuli/index.html) and were shown in white against a black background at a size of ∼ 250 × 520 px. The 16 blocks were divided randomly into 4 experimental conditions, constructed from combining the manipulations of phase scrambling (100% scrambled *vs*. non-scrambled) and inversion (inverted *vs*. upright). Phase scrambling was implemented by randomly changing the phase of each dot to another phase between 0 and 360°. To mitigate potential habituation effects, each repetition of the same condition showed a different stimulus, namely a male or female walker, facing left or right. Furthermore, to control participants’ eye gaze and their attentional focus on the screen, the central dot of the walker was colored in grey and served as a fixation cross. Participants were asked to focus on this dot and to press the spacebar every time it turned red (400 ms), which happened two to four times per block. Detection accuracy was high across all conditions (i.e., 97-98%).

### Preprocessing

EEG was recorded from 64 Ag/AgCI (active) electrodes using an ActiCHamp amplifier and BrainVisionRecorder software (version 1.21.0402, Brain Products, Gilching, Germany) at a sampling rate of 1000 Hz. Electrode positions were based on the 10%-system, except for two electrodes (TP9 and TP10), which were placed at OI1h and OI2h according to the 5%-system to have better coverage of posterior scalp sites. Fpz was used as ground electrode and Fz was used as online reference. Horizontal eye movements were recorded with the FT9 and FT10 electrodes embedded in the EEG cap and vertical eye movements with two additional bipolar AG/AgCI sintered ring electrodes placed above and below the left eye.

Off-line processing of the EEG signal was done in Letswave 6 (www.letswave.org) according to the following steps. First, the raw data were band-pass filtered using a fourth-order Butterworth filter with 0.1 Hz and 100 Hz as cut-off values. Next, the filtered data were segmented according to the 4 experimental conditions (-2 to 54 sec), before an independent component analysis (ICA) was applied on the merged segmented data to remove ocular artefacts. Following ICA, faulty or excessively noisy electrodes were interpolated (2% on average, never more than 10%) and the data were re-referenced to the average signal across all electrodes. Next, the re-referenced epochs were cropped into epochs running from the end of the fade-in to the start of the fade-out period to ensure that epoch length was a multiple of the presentation rate. Finally, conditions were averaged and a Fast Fourier Transform was applied to transform the data to normalized (divided by N/2) amplitudes (μV) in the frequency domain.

### Data Analysis

Frequency tagging elicits not only responses at the tagged frequency (F) but also at harmonics of that frequency (2F, 3F, …). Given that the brain response is spread out across these different frequencies (Norcia et al., 2015), it is best quantified by summing the baseline-subtracted amplitudes across all relevant harmonics (Retter et al., 2021; Retter & Rossion, 2016). Therefore, as preregistered, we first determined the number of harmonics to include by (i) taking the grand-averaged amplitudes across participants, conditions, and electrodes, (ii) calculating a z-score for each frequency bin using the 10 surrounding bins on each side as baseline, excluding directly adjacent bins, and (iii) identifying the harmonics with a z-score > 2.32 (i.e., p < .01; for a similar approach, see Cracco et al., 2021; Retter & Rossion, 2016). This procedure identified 3 relevant harmonics: 2.4 Hz (F), 4.8 Hz (2F), and 7.2 Hz (3F). For each of these 3 harmonics, we then calculated the baseline-subtracted amplitudes for each participant, condition, and electrode using the same baseline as for the z-scores and these amplitudes were summed to quantify the brain response (Retter et al., 2021; Retter & Rossion, 2016).

In addition to a response at 2.4 Hz, the grand-averaged amplitude spectrum also revealed a response at 1.2 Hz. This latter frequency is the frequency at which each individual dot repeated its trajectory. In the non-scrambled condition, this means that it is the frequency at which the walker took a step with the same foot. Although we had no a-priori predictions regarding this response, we nevertheless analyzed it in a preregistered secondary analysis, using the same procedure as above, but excluding those harmonics that overlapped with the 2.4 Hz response, as recommended by Retter et al. (2021). The analysis of the 1.2 Hz response revealed 2 harmonics with z > 2.32, at 3.6 Hz (2F) and 6 Hz (3F). Although not exceeding the pre-defined z-threshold, a smaller peak was also visible at 1.2 Hz (F). Given that noise is typically higher at low frequencies, it is possible that this reduced peak at 1.2 Hz did not reflect a weak response, but rather high noise. We therefore decided to also include the 1.2 Hz amplitudes in the main analysis, but found that excluding these amplitudes did not change the results.

To determine the electrodes of interest, we used a preregistered collapsed localizer approach using the noise-subtracted amplitudes (Luck & Gaspelin, 2017). That is, we identified the relevant electrodes from the averaged topography across participants, conditions, and both responses (i.e., 1.2 Hz and 2.4 Hz). The averaged topography revealed widespread activity across all occipital, parieto-occipital, and parietal electrodes, with a slight right-sided lateralization (Figure 2). Given that such lateralization matches previous work on biological motion perception (e.g., Grossman et al., 2000; Jokisch et al., 2005), we decided to formally include laterality in the analysis. More specifically, we divided the activated scalp area into two clusters comprising all O, PO, and P electrodes, except for the five electrodes on or around the midline (Oz, OI1h, OI2h, POz, and Pz), and tested for differences between these two clusters by adding laterality (left *vs*. right) as a factor to the design. Note, however, that including the five excluded electrodes did not change the results (see Supplementary Analysis).

**Figure 2.**
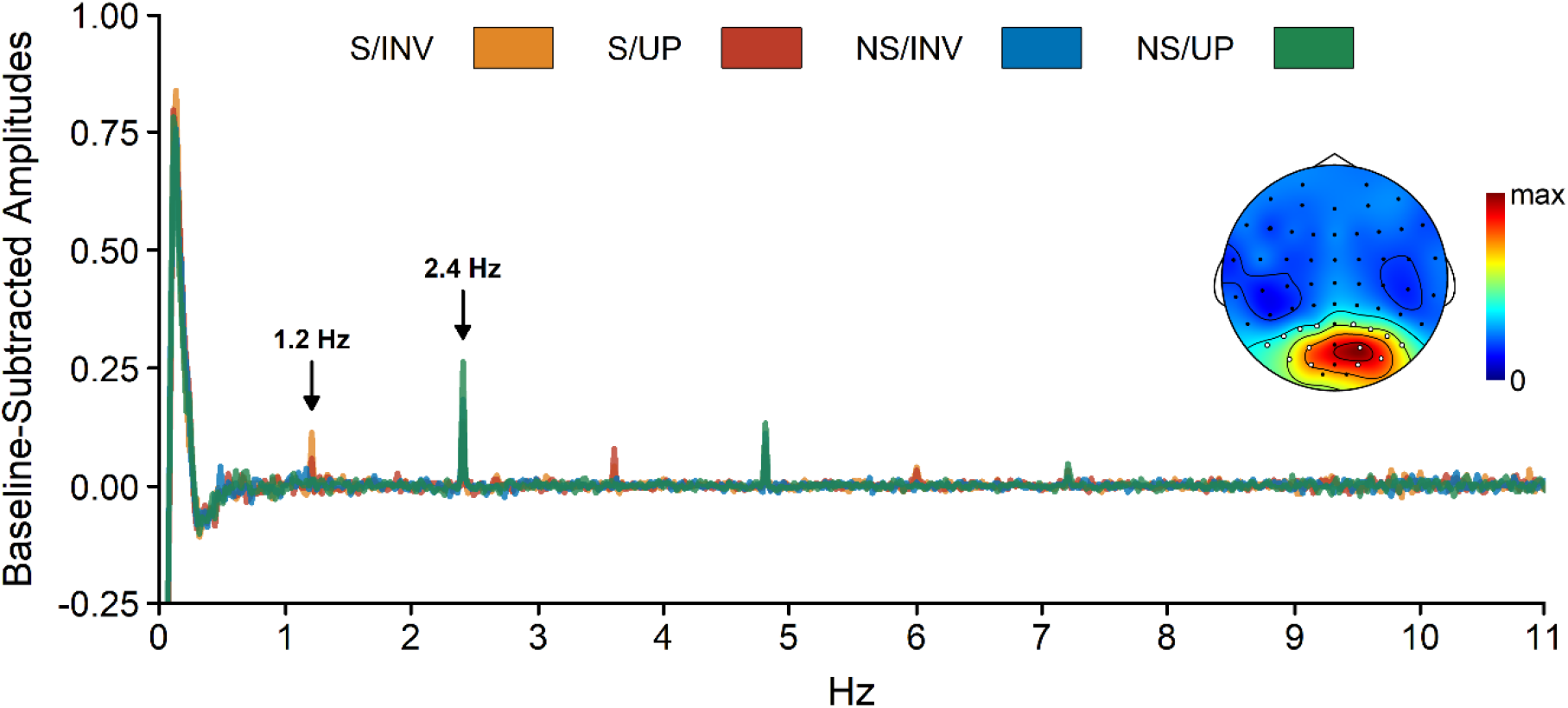
Collapsed Topography and Spectrum Plot of the Baseline-Subtracted Amplitudes *Note*. The collapsed topography shows the baseline-subtracted amplitudes across participants, conditions, and both brain responses. Electrodes included in the analysis are marked in white. The topography is scaled from 0 to the maximum value across all electrodes (i.e., 0.32 μV). The spectrum plot shows the baseline-subtracted amplitudes for the included electrodes, separately for the four conditions. S = scrambled, NS = non-scrambled, INV = inverted, UP = upright.

Using the above electrodes, we conducted separate repeated measures ANOVAs for the 2.4 Hz and 1.2 Hz responses using scrambling (scrambled *vs*. non-scrambled), orientation (inverted *vs*. upright), and laterality (left *vs*. right) as within-subject factors. Furthermore, as a preregistered exploratory analysis, we also correlated the brain responses at both frequencies with the collected AQ scores. Note, however, that this latter analysis is likely underpowered.

## Results

### 2.4 Hz Analysis

Amplitudes at 2.4 Hz (Figure 3) measure brain responses coupled to the walking cycle. Analyzing these responses revealed a main effect of scrambling, *F*(1, 29) = 45.64, *p* < .001, *d*_z_ = 1.23, with stronger responses in the non-scrambled than in the scrambled condition, a main effect of orientation, *F*(1, 29) = 5.36, *p* = .028, *d*_z_ = 0.42, with stronger responses in the upright than in the inverted condition, and a main effect of laterality, *F*(1, 29) = 12.30, *p* = .002, *d*_z_ = 0.64, with stronger responses across right-sided than across left-sided electrodes. In addition, we found an interaction between scrambling and orientation, *F*(1, 29) = 8.50, *p* = .007, *d*_z_ = 0.53, indicating that there was an effect of inversion in the non-scrambled condition, *t*(29) = 3.08, *p* = .005, *d*_z_ = 0.56, but not in the scrambled condition, *t*(29) = 0.47, *p* = .640, *d*_z_ = 0.08. None of the remaining effects reached statistical significance, all *F* ≤ 2.04, all *p* ≥ .164. Finally, exploratory Spearman correlations revealed no significant assocations between the AQ total score and the main effect of scrambling, *r*_s_ = 0.10, *p* = .609, the main effect of orientation, *r*_s_ = 0.28, *p* = .134, or the scrambling x orientation interaction effect, *r*_s_ = 0.17, *p* = .361.

**Figure 3.**
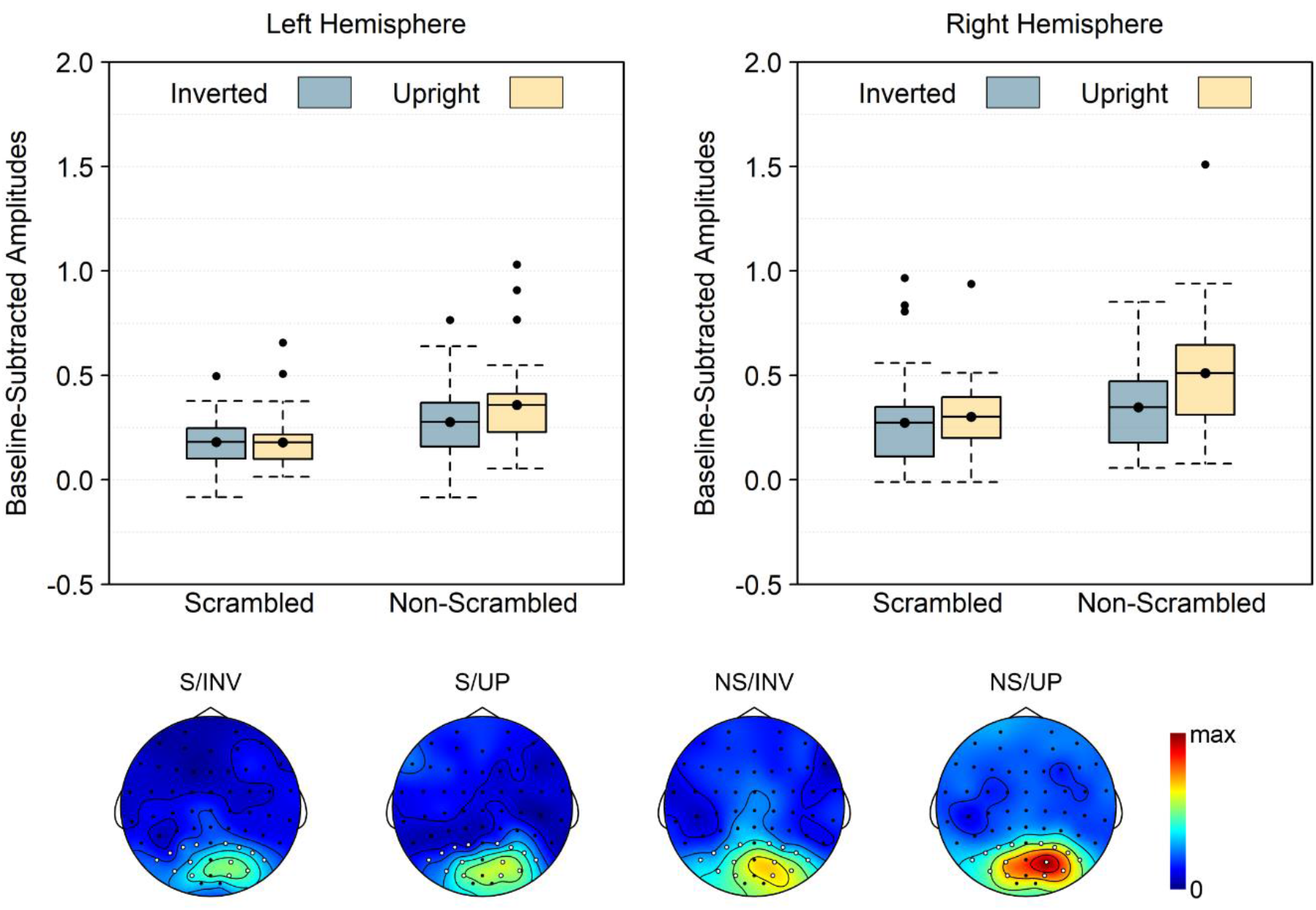
Baseline-Subtracted Amplitudes at 2.4 Hz and Harmonics *Note*. Boxplots show the mean instead of median to match the statistical analysis. Note that 0 is the baseline and that values below 0 necessarily reflect noise. Topographies are plotted on a scale from 0 to the maximum across all 4 conditions (i.e., 0.74 μV), with the included electrodes marked in white. S = scrambled, NS = non-scrambled, INV = inverted, UP = upright.

### 1.2 Hz Analysis

Amplitudes at 1.2 Hz (Figure 4) measure brain responses coupled to the movement cycles of the individual dot. Analyzing these responses revealed an inversed main effect of scrambling, *F*(1, 29) = 66.26, *p* < .001, *d*_z_ = 1.49, with stronger responses in the scrambled than in the non-scrambled condition. Strikingly, *t* tests comparing the 1.2 Hz response to baseline revealed that it was *only* present in the two scrambled conditions, both *t* ≥ 7.51, both *p*_one-tailed_ < .001, but not in the two non-scrambled conditions, both *t* ≤ 1.20, both *p*_one-tailed_ ≥ .120 (see also Figure 2). In contrast to the 2.4 Hz response, the main effects of orientation, *F*(1, 29) = 0.47, *p* = .499, *d*_z_ = 0.13, or laterality, *F*(1, 29) = 1.30, *p* = .265, *d*_z_ = 0.21, were not significant, and neither was the scrambling x orientation interaction, *F*(1, 29) = 0.32, *p* = .577, *d*_z_ = 0.10. The laterality x scrambling interaction, however, did reach significance, *F*(1, 29) = 8.90, *p* = .006, *d*_z_ = 0.55, indicating that the effect of scrambling was stronger across right, *t*(29) = 7.73, *p* < .001, *d*_z_ = 1.41, than across left electrodes, *t*(29) = 6.10, *p* < .001, *d*_z_ = 1.11. None of the other effects reached significance, all *F* ≤ 1.04, all *p* ≥ .316. Exploratory Spearman correlations revealed no significant relation between the AQ total score and the main effect of scrambling, *r*_s_ = 0.04, *p* = .825, or the laterality x scrambling interaction effect, *r*_s_ = -0.01, *p* = .965.

**Figure 4.**
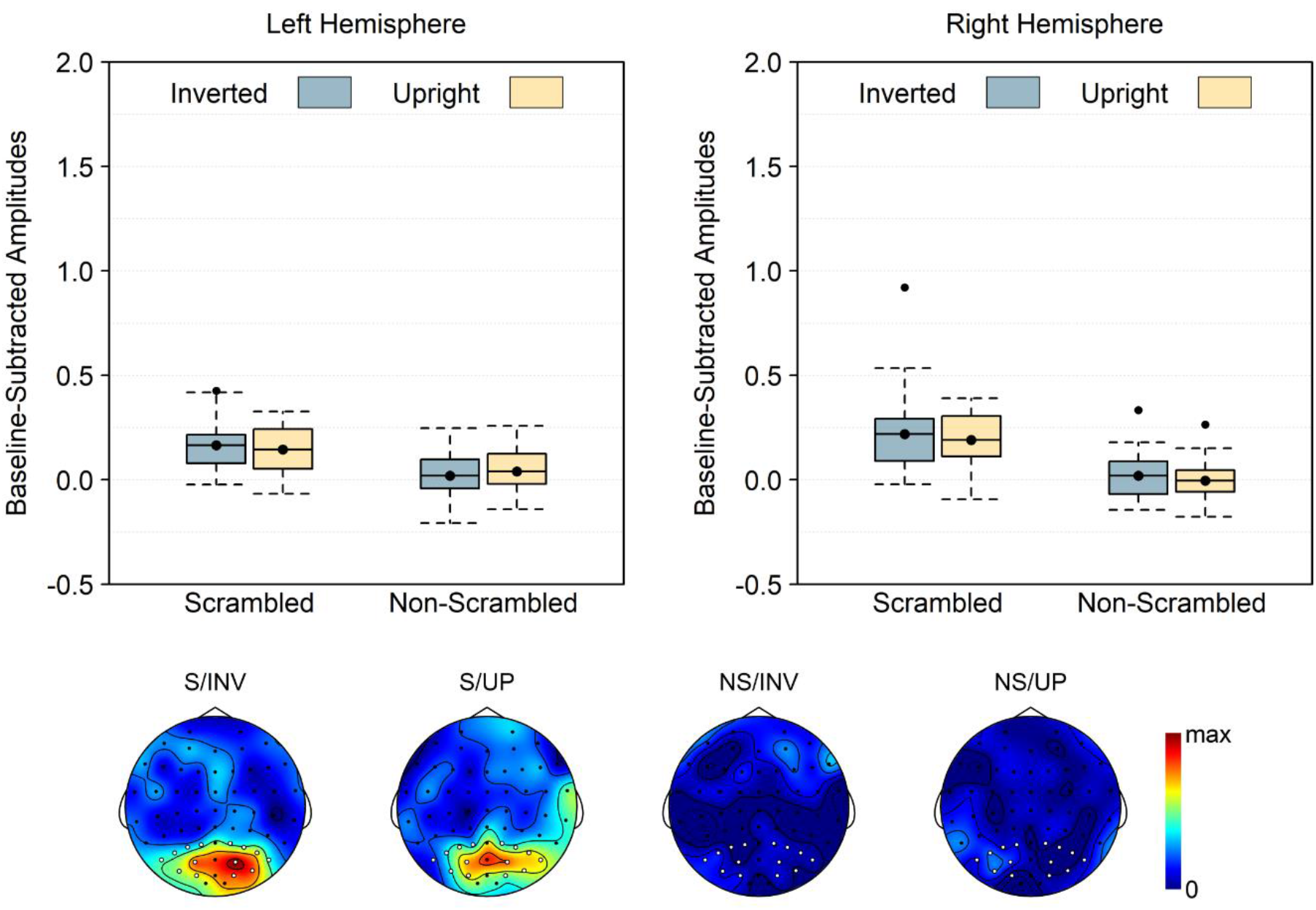
Baseline-Subtracted Amplitudes at 1.2 Hz and Harmonics *Note*. Boxplots show the mean instead of median to match the statistical analysis. Note that 0 is the baseline and that values below 0 necessarily reflect noise. Topographies are plotted on a scale from 0 to the maximum across all 4 conditions (i.e., 0.34 μV), with the included electrodes marked in white. S = scrambled, NS = non-scrambled, INV = inverted, UP = upright.

## Discussion

Separating movement processing from other processes also activated during movement perception is a key challenge in the study of biological movement perception. Here, we did so by presenting a point-light figure walking at a pace of 2.4 Hz and measuring brain responses recurring at exactly that pace, thereby naturally disentangling movement processing from concomitant secondary processes not coupled to the movement itself (e.g., extracting social traits). Validating our approach, we found reliable brain responses at the frequency of walking that were reduced by two manipulations well-known to disrupt movement processing: phase scrambling (e.g., Beintema et al., 2006; Troje & Westhoff, 2006) and stimulus inversion (e.g., Bertenthal & Pinto, 1994; Pavlova et al., 2017; White et al., 2014).

Coupling brain responses to the repetition of a cyclical movement has three key advantages over existing methods. First, in contrast to previous fMRI and ERP studies, movement tagging discards all brain processes not coupled exactly to the temporal profile of the movement, thereby providing an objective signature of visual movement processing. Second, it provides not only an objective but also an ecologically valid signature of biological movement perception, as it captures the online processing of observed movements as they occur, rather than the processing of brief and sometimes incomplete movements, artificially divided into different trials. Finally, although not predicted a-priori, movement tagging appears to naturally disentangle two ways of processing point-light figures: they can be processed globally, as a moving agent, or locally, as a set of moving dots (Chang & Troje, 2009b). In our task, global processing was captured by the brain response at 2.4 Hz, the walking pace. Indeed, supporting this view, amplitudes at 2.4 Hz were reduced when scrambling perturbed the global movement percept. However, in addition to the predicted response at 2.4 Hz, we also found a response at 1.2 Hz, coupled to the individual dot cycles. In contrast to the 2.4 Hz response, this 1.2 Hz response was not decreased but increased by scrambling. Thus, our results indicate that movement tagging can disentangle global (2.4 Hz) and local (1.2 Hz) movement within the same stimulus and that disrupting processing at one level increases processing at the other, consistent with previous evidence of interactions between local and global processes in biological motion perception (e.g., Hirai et al., 2011; Wang et al., 2010).

The idea that the 2.4 Hz response was specific to global movement processing is further supported by the finding that it was susceptible to inversion only when the stimulus was not scrambled. Indeed, previous research has shown that inversion perturbs not only global movement processing but also local motion processing of the feet (Chang & Troje, 2009a; Troje & Westhoff, 2006). This, in turn, causes inversion to have similar effects on scrambled and non-scrambled walkers (Troje & Westhoff, 2006). The fact that we did *not* find an inversion effect for the scrambled walker in the current study therefore indicates that the measured brain responses at 2.4 Hz were a relatively pure measure of global movement perception.

Going one step further, the pattern of results at 1.2 Hz suggests that inversion likely did not influence local motion processing at all in the current task. Indeed, whereas scrambling increased the 1.2 Hz response, inversion did not. In contrast to Troje and Westhoff (2006), this suggests that inversion had a rather specific influence on global movement perception. A likely reason for this discrepancy is that we presented the walker for an extended duration of ∼ 1 minute without task instructions, while Troje and Westhoff (2006) presented their stimuli for no longer than a few seconds and asked participants to identify the direction in which the figure was moving. Research has shown that such direction discrimination tasks can be solved easily based on local motion cues alone (e.g., Chang & Troje, 2009b; Troje, 2013). Given that local cues can be processed quickly in pre-cortical brain areas (Hirai & Senju, 2020), they likely dominate fast-paced direction discrimination tasks (e.g., Hirai, Saunders, et al., 2011). Showing the same stimulus for an extended period, on the other hand, may instead trigger a more global processing style. Indeed, it seems highly unlikely that participants in the current study did not realize that the inverted walker was still a walker. Hence, a plausible explanation of our findings is that both upright and inverted walkers were processed globally but that inverted walkers were processed less efficiently because they did not map neatly onto existing templates (Giese & Poggio, 2003; Lange & Lappe, 2006, 2007).

Nevertheless, despite the above, any comparison of the responses at 1.2 and 2.4 Hz must take into account that they are harmonically related. In other words, an important question is how we can be sure that amplitudes at those two frequencies were not just two components of the same response, rather than two different responses. Two points support the latter option. First, a response at 1.2 Hz was *only* visible when the point-light walker was scrambled, making it very unlikely that the response at 2.4 Hz was merely a harmonic of the slower 1.2 Hz response in the non-scrambled conditions. Yet, it remains possible that the 2.4 Hz response was simply a 1.2 Hz harmonic in the scrambled conditions, where we did see a reliable 1.2 Hz response. However, this again seems unlikely, considering that scrambling had opposite effects at 1.2 and 2.4 Hz. Together, these findings clearly indicate that despite their harmonic relationship, amplitudes at 1.2 and 2.4 Hz tagged distinct processes (see also Cracco et al., 2021).

In sum, the movement tagging method developed here not only directly and objectively captures the online processing of biological movements, but also naturally disentangles local and global movement processing. The latter is important, because the two types of movement processing are known to rely on different mechanisms (e.g., Chang & Troje, 2009b). Indeed, according to a recent model, there are two main mechanisms involved in the perception of walking: a ‘step detection mechanism’ and an ‘action body evaluator’ (Hirai & Senju, 2020; see also Neri, 2009). The first mechanism is thought to develop early in life and relies on local stimulus information such as the movement of the feet. By guiding attention to biological motion, this mechanism then gradually leads to the development of the second mechanism, involved in processing global body actions. Testing the predictions of this two-process model critically requires a method that is able to disentangle both processes. Our results suggest that movement tagging may be able to do this without having to develop ecologically invalid, artificial stimuli that are manipulated in specific ways to contain only local or only global stimulus information (e.g., Chang et al., 2018; Chang & Troje, 2009b).

Distinguishing global from local movement processing may also be important to elucidate the biological motion perception anomalies that have been reported in autism (Federici et al., 2020; Todorova et al., 2019; Van der Hallen et al., 2019). Indeed, while some studies have found such anomalies (e.g., Nackaerts et al., 2012), other studies failed to find any difference between individuals with and without autism (e.g., Edey et al., 2019). Given that perceptual differences in autism mostly involve global processes (Happé & Frith, 2006; Van der Hallen et al., 2015, 2019), it seems likely that one reason for this inconsistency is that local and global processes are often confounded in biological motion tasks (Chang & Troje, 2009b). By teasing those two processes apart, movement tagging could thus help explain exactly which aspects of biological motion perception differ between individuals with and without autism.

However, while our results clearly show that movement tagging can be used to objectively measure movement tagging, more work is still needed to determine its boundary conditions. For example, the current study used only one type of action, walking. Although walking is by far the most commonly used action in the literature and forms the basis of existing theoretical models (Giese & Poggio, 2003; Hirai & Senju, 2020; Lange & Lappe, 2006), further research will have to investigate whether movement tagging works with other movements as well. It seems likely that it will, however, as long as the movement is cyclical (for a database of cyclical actions, see Vanrie & Verfaillie, 2004). In addition, the current study investigated only one type of scrambling, temporal scrambling, in which the phase but not the position of the different dots is scrambled. It, thus, remains an open question whether spatial scrambling has similar effects. Yet this again seems likely, given that previous research has shown, using a different approach, that spatial and temporal scrambling have very similar effects on biological motion perception (e.g., Troje & Westhoff, 2006). Finally, the current study used only one presentation frequency (2.4 Hz) and hence it remains unclear to what extent similar results would be obtained with other frequencies. That said, previous research on, for example, face perception has shown that frequency tagging usually yields comparable responses across a wide range of frequencies (Retter & Rossion, 2016).

To conclude, the current study used frequency tagging to objectively measure biological motion perception. In addition to separating movement processing from other, concomitant processing, this has the crucial advantage that it can disentangle local and global movement processing at different frequencies of the same brain signal. These results open up important new avenues for studying the visual processes that contribute to biological motion perception.

## Supplementary Material

### Analysis Including Middle Posterior Electrodes

**Figure S1.**
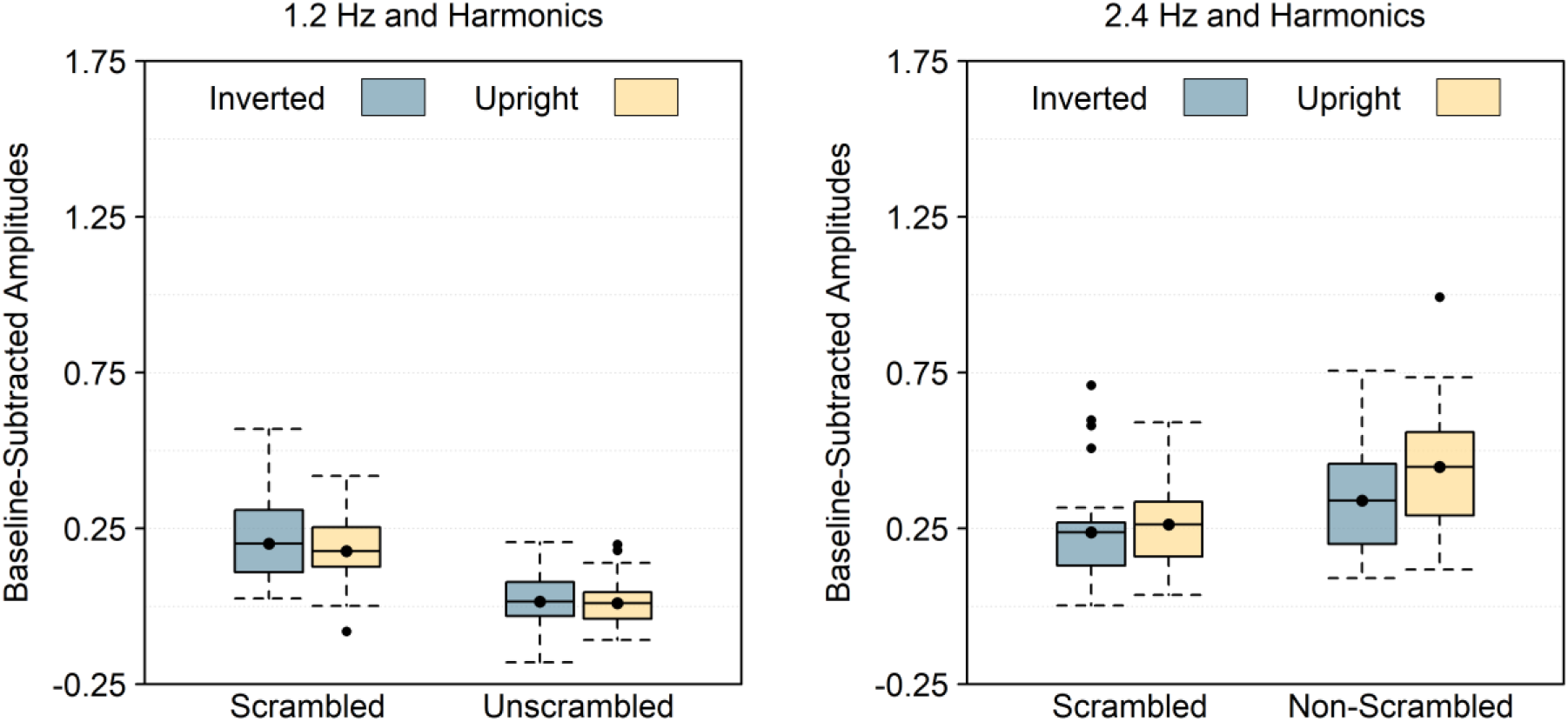
Baseline-Subtracted Amplitudes at 1.2 Hz and 2.4 Hz and Harmonics *Note*. Boxplots show the mean instead of median to match the statistical analysis. The following electrodes are included in addition to the electrodes included in the main analysis: OI1h, OI2h, Oz, POz, and Pz. Note that 0 is the baseline and that values below 0 necessarily reflect noise.

### 2.4 Hz Analysis

When the five electrodes on or around the midline were included, the 2.4 Hz analysis (Figure S1) revealed a main effect of scrambling, *F*(1, 29) = 47.17, *p* < .001, *d*_z_ = 1.25, with stronger responses in the non-scrambled than in the scrambled condition, a main effect of orientation, *F*(1, 29) = 5.22, *p* = .030, *d*_z_ = 0.42, with stronger responses in the upright than in the inverted condition, and a scrambling x orientation interaction, *F*(1, 29) = 5.52, *p* = .026, *d*_z_ = 0.43, indicating that the orientation effect was present in the non-scrambled condition, *t*(29) = 2.68, *p* = .012, *d*_z_ = 0.49, but not in the scrambled condition, *t*(29) = 0.95, *p* = .352, *d*_z_ = 0.17.

### 1.2 Hz Analysis

When the five electrodes on or around the midline were included, the 2.4 Hz analysis (Figure S1) revealed a main effect of scrambling, *F*(1, 29) = 69.17, *p* < .001, *d*_z_ = 1.52, with stronger responses in the scrambled than in the non-scrambled condition, but no main effect of orientation, *F*(1, 29) = 0.77, *p* = .389, *d*_z_ = 0.16, nor a scrambling x orientation interaction, *F*(1, 29) = 0.26, *p* = .618, *d*_z_ = 0.09.

